# Bronchoalveolar lavage affects thorax computed tomography of healthy and SARS-CoV-2 infected rhesus macaques (*Macaca mulatta*)

**DOI:** 10.1101/2021.02.04.429761

**Authors:** Annemiek Maaskant, Lisette Meijer, Jaco Bakker, Leo van Geest, Dian G.M. Zijlmans, Jan A.M. Langermans, Ernst J. Verschoor, Marieke A. Stammes

## Abstract

Medical imaging as method to assess the longitudinal process of a SARS-CoV-2 infection in non-human primates is commonly used in research settings. Bronchoalveolar lavage (BAL) is also regularly used to determine the local virus production and immune effects of SARS-CoV-2 in the lower respiratory tract. However, the potential interference of those two diagnostic modalities with each other is unknown in non-human primates. The current study investigated the effect and duration of BAL on computed tomography (CT) in both healthy and experimentally SARS-CoV-2-infected female rhesus macaques (*Macaca mulatta*). In addition, the effect of subsequent BALs was reviewed. Thorax CTs and BALs were obtained from four healthy animals and 11 experimentally SARS-CoV-2-infected animals. From all animals, CTs were obtained just before BAL, and 24 hours post-BAL. Additionally, from the healthy animals, CTs immediately after and four hours post-BAL were obtained. Thorax CTs were evaluated for alterations in lung density, measured in Hounsfield units, and a visual semi-quantitative scoring system. An increase in the lung density was observed on the immediately post-BAL CT but resolved within 24 hours in the healthy animals. In the infected animals, a significant difference in both the lung density and CT score was still found 24 hours after BAL. Furthermore, the differences between timepoints in CT score were increased for the second BAL. These results indicate that the effect of BAL on infected lungs is not completed within the first 24 hours. Therefore, it is of importance to acknowledge the interference between BAL and CT in rhesus macaques.

## Introduction

Infection with Severe Acute Respiratory Syndrome Coronavirus 2 (SARS-CoV-2) and the development of coronavirus 2019 (COVID-19) was first described in Wuhan, Hubei Province, China at the end of 2019 (1). Since then, SARS-CoV-2 spread rapidly across the globe, leading the World Health Organization to officially declare COVID-19 a Public Health Emergency of International Concern by the end of January 2020.

Animal models to investigate the infection progress, disease development, and to evaluate prophylactic and therapeutic treatment options are important to answer several research questions. Due to the close resemblance of their physiology and immune system to that of man, non-human primates (NHPs) play a pivotal role in those aforementioned research topics (2–4). Multiple studies compared several NHP models to identify their suitability for COVID-19 research, and demonstrated that Old World monkeys, like macaques, are susceptible to SARS-CoV-2 infection and develop a mild-to-moderate form of disease (5–8).

Computed tomography (CT) of the thorax and bronchoalveolar lavage (BAL) are diagnostic procedures widely used in clinical pulmonary research and veterinary medicine (9–12). These techniques are also used in COVID-19 research in NHPs to gain more information and to create a bridge between the human and NHP data (6, 13, 14). However, to the authors knowledge, no literature concerning the effect of BAL on CT is available regarding NHPs.

For SARS-CoV-2, the infection is predominantly found in the respiratory tract. To determine infection and viral load, nasopharyngeal and tracheal swabs are used to sample the upper respiratory tract. With a BAL, samples from the lower respiratory tract can be obtained (12, 15). BAL fluid can be used both as diagnostic tool and for detailed analysis of the infection. This includes a variety of analytical tests such as cell counts and differential, cytopathologic analysis, and cultures in addition to specific molecular and immunologic diagnostic tests (9). In COVID-19 research, it is predominantly used to prove the presence of virus locally in the lungs. By combining swabs with a BAL, the majority of the respiratory tract can be screened. Nevertheless, the utility of BAL in COVID-19 research or as diagnostic procedure, in general, is arguable. A potential drawback of the use of BAL is the possibility of artificial dissemination of SARS-CoV-2 and the impact on the readout of thorax CTs (13). In addition, Geri *et al*. suggests that when the swabs of the upper respiratory tract and thorax CTs are negative it is likely that the BAL is negative as well (16). Others state that BAL has a place in diagnosis of COVID-19 besides swabs and imaging or that, in vaccine testing, BAL is more robust and specific to demonstrate reduction in viral load than in nasal swabs and more specific than the lesions found on thorax CTs (17–19).

The basis for successful execution of our COVID-19 infection studies is the use of an optimally-characterized macaque infection model. We investigated the effect of BAL on thorax CTs of rhesus macaques. Additionally, we determined the timeframe of possible interference as during BAL a part of the instilled saline is left unrecovered in the lungs (12). Studies performed in dogs and humans demonstrated that retained saline is correlated with an increase in lung opacity on immediate post-BAL CTs (12, 15). By this the retained saline can mask COVID-19 associated lesions on CT as one of the main groups of those lesions are ground glass opacities (GGOs) which are also characterized by a slight increase in lung opacity (13, 20).

The aim of this study is to evaluate the time effects and potential impact of subsequent BALs on the visualization and quantification of the lungs with CT in healthy and SARS-CoV-2 infected rhesus macaques to appraise possible differences during a viral infection.

## Materials and Methods

### Animals & Procedures

The study consisted of two groups of rhesus macaques; a non-infected group and a SARS-CoV-2 infected group. The study was conducted at the Biomedical Primate Research Centre (BPRC, Rijswijk, The Netherlands). For the non-infected group, four healthy rhesus macaques were included. Three BAL samples were taken to evaluate potential deleterious effects on the lung by using various thorax CTs. For the infected group 11 macaques were analyzed, derived from three different COVID-19 studies. These animals were selected out of those studies of which a CT was obtained before and 24 hours after a BAL during the active phase of the infection. In addition, in four infected macaques an additional BAL at day 2 and an additional CT immediately post-BAL was obtained.

As a timeline, for the non-infected animals three BALs were performed; on Day 0, 2 and 6 respectively BAL 1, 2 and 3. With each BAL four CTs are obtained; pre-BAL, post-BAL, four hours post-BAL and 24 hours post-BAL. For the infected animals one BAL is performed coupled with two CTs: pre-BAL and 24 hours post-BAL. In addition, four of the infected animals underwent a subsequent BAL at day 2, 48 hours after the first BAL with a pre-BAL CT, post-BAL CT and 24 hours post-BAL CT.

All animals were of Indian origin, female rhesus macaques (*Macaca mulatta*), purpose bred at the BPRC. During the study the animals were socially housed in pairs. All animals were between 7 and 16 years old and weighed between 6.2 to 11.5 kg. Cages for each pair of animals measured 1 x 2 x 2 meters and were divided into different compartments that were freely accessible to allow animals the choice where to sit. All cages were provided with bedding to allow foraging, environmental enrichment such as fire hoses, mirrors and toys. Daily, all animals also received additional food enrichment items. All procedures, husbandry, and housing performed in this study were in accordance with the Dutch laws on animal experimentation and the regulations for animal handling as described in EU Directive 63/2010.

BPRC is accredited by the Association for Assessment and Accreditation of Laboratory Animal Care (AAALAC) International. The study was performed under a project license issued by the Competent Authorities (Central Committee for Animal experiments, license no. AVD5020020209404). Before the start of the study, approval was obtained, by the institutional animal welfare body (CCD 028 Evaluation of vaccines and antiviral compounds against emerging coronavirus infections).

The procedures were all performed in the morning. All macaques were fasted overnight before the procedure while water was freely available throughout. Animals were anesthetized in their home cage with ketamin (10 mg/kg, ketamine hydrochloride, Ketamin 10%; Alfasan Nederland BV, Woerden, the Netherlands) combined with medetomidine hydrochloride (0.05 mg/kg, Sedastart; AST Farma B.V., Oudewater, the Netherlands) intramuscular (IM) and were transferred to the CT-room. At the end of the procedures, when the macaques returned to their home cage, atipamezole (0.25 mg/kg, Sedastop; AST Farma B.V., Oudewater, the Netherlands) was administered IM. In addition, during every sedation bodyweight was measured.

### Bronchoalveolar lavage

Fiber-optic bronchoscopy (FOB) with BAL targeting the left lung was performed at various timepoints. The procedure was executed as described before (21). Three volumes of 20 ml of prewarmed sterile 0.9% saline solution (NaCl 0.9%; B. Braun medical B.V., Oss, the Netherlands) were consecutively instilled in the left lung and subsequently aspirated. In all cases at least 85% of the fluid was recovered. After BAL the animals were transported back to the CT-room, located next to the BAL-room, were the CT was immediately obtained. For the time between the subsequent scans, the animals were transferred back to their home cage.

### Computed tomography

Non-invasive, free breathing, CT data were acquired on several timepoints but also always pre-infection using a MultiScan Large Field of View Extreme Resolution Research Imager (LFER) 150 PET-CT (Mediso Medical Imaging Systems Ltd., Budapest, Hungary). The macaques were positioned head first supine (HFS) with the arms up and fixated in a vacuum pillow. A single CT of the thorax takes 35 seconds by which respiratory motion is inevitable. To mitigate the impact of respiratory motion and improve the image quality, respiratory gating was applied. The respiratory amplitude was detected with a gating pad placed next to the umbilicus. For the final reconstruction, the inspiration phases were exclusively used and manually selected (22).

For five of the SARS-CoV-2 infected animals, the 24 hours post-BAL CT was part of a PET-CT procedure. This procedure is described elsewhere (23).

### SARS-CoV-2 infection

A group of 11 animals was exposed to a dose of 1 x 10^5^ TCID_50_ SARS-CoV-2, diluted in phosphate buffered saline (PBS). The virus was inoculated via a combination of the intratracheal route and intranasal route. During the infection period the animals were checked twice daily by the animal caretakers and scored for clinical symptoms. All animals used in this project had a confirmed SARS-CoV-2 infection as determined by multiple positive RT-PCR scores. RT-PCR scores were determined as described elsewhere (24, 25). The clinical and pathological data will be described separately.

## Data analysis

### CT scoring

A semi-quantitative scoring system for thorax CT evaluation was used to estimate SARS-CoV-2-induced lung disease (7, 13). Quantification of the opacities on the CTs was performed independently by two experienced imaging scientists based on the sum of the lobar scores. The degree of involvement in each zone was scored as: 0 for no involvement, 1 for <5%, 2 for 5-24%, 3 for 25-49%, 4 for 50-74% and 5 for >=75% involvement. An additional increase or decrease of 0.5 was used to indicate alterations in CT density of the lesions. By using this scoring system, a maximum score of 35 could be reached for the combined lobes per time point.

### Lung density

In each CT, in two representative planes, in the coronal direction, regions of interest (ROIs) were manually drawn over the left and right lung with a slice thickness of 12 mm using Vivoquant v4.5 (InVicro, Boston, USA). The first plane was defined in the center of the carina. The second plane was drawn more ventral at a predefined distance of 50 mm from the first plane ending up roughly in the center of the heart. Of those ROIs a histogram was generated representing the voxel intensity, measured in Hounsfield units (HUs) ranging from −1000 till 0. HUs are linearly correlated with physical density and in this way alterations in the lung density pattern after BAL could be visualized. For analyses the histogram was divided into three parts; −1000/−600, −600/−400, and −400/0, which were defined as low density (LD), medium density (MD) and high density (HD) respectively. The LD was defined as healthy lung based on the average value of the non-infected animals. The HD was characterized as non-healthy lung tissue and the MD as in between, in this range the differences between scans were also captured due to the normal existing differences in respiratory amplitude and pattern. Afterwards absolute values were converted to relative values to compensate for the difference in lung volume both between scans and between animals and calculated as area under the curve (AUC).

### Statistical analysis

Statistical tests were performed with SPSS statistics v26 and GraphPad prism v8.4.2. After testing for normal distribution, the t-test was used to compare the results obtained from the histograms and the Wilcoxon test for CT scores. For correlation analysis, respectively a Pearson and Spearman correlation were applied. The level of significance was α=0.05, all tests were performed two-sided and where possible paired.

## Results

Before comparing the different groups and timepoints with each other, the entire group of animals was examined for possible confounding factors in relation to age and weight. In correlation with both visual and quantitative assessment no confounding factors were found for these parameters.

### Visual assessment non-infected group

In all non-infected animals, an increase in density was visible in the lavaged lung around the opening of the main bronchus at the CT performed immediately post-BAL. The increased density was more focused in either the middle or the lower lobe. Overall the spread of the BAL fluid was more in the dorsal direction of the lungs than the ventral direction. At the scan obtained four hours post-BAL, the majority of these alterations observed immediately post-BAL were resolved. In addition, the density of the lesions was decreased. At the scan obtained 24 hours after the BAL, the lungs had visually returned to baseline values obtained just before BAL. This pattern was similar in all BALs. The timeline and representative The individual scores pre-/post-BAL of the CT images on the different timepoints are summarized in Table 1. Due to a technical problem, one of the non-infected animals was lavaged in the right lung instead of the left. This animal was excluded from further data analysis.

**Table 1:** overview of the CT scores of the left (le) and right (ri) lung of non-infected rhesus macaques (RM) at all timepoints

| RM | BAL | CT score pre |  | CT score post |  | CT score 4h post |  | CT score 24h post |  |
| --- | --- | --- | --- | --- | --- | --- | --- | --- | --- |
|  |  | le | ri | le | ri | le | ri | le | ri |
| 1 | 1 | 1 | 0 | 9 | 0 | 0 | 0 | 1 | 0 |
|  | 2 | 2 | 0 | 11 | 4 | 1 | 0 | 0 | 0 |
|  | 3 | 2 | 0 | 9 | 0 | 1 | 0 | 1 | 0 |
| 2 | 1 | 0 | 0 | 9 | 0 | 2 | 0 | 1 | 0 |
|  | 2 | 5 | 0 | 8 | 0 | 5 | 0 | 1 | 2 |
|  | 3 | 0 | 0 | 9 | 0 | 1 | 0 | 1 | 0 |
| 3 | 1 | 4 | 0 | 10 | 0 | 3 | 0 | 5 | 0 |
|  | 2 | 6 | 0 | 8 | 0 | 3 | 0 | 1 | 6 |
|  | 3 | 5 | 0 | 8 | 1 | 2 | 0 | 4 | 0 |

The average difference in CT score for the left lung between the pre-BAL CT and the CT obtained after 24 hours was 0.2, confirming that at 24 hours post-BAL the appearance of the lungs is comparable to the situation pre-BAL. Nevertheless, when comparing the subsequent BALs with each other, the CT score pre-BAL 2 and four hours post-BAL 2 were higher compared to the related time points of BAL 1 and BAL 3.

Post-BAL the left lung scored between 8-11, a significantly higher score compared to all pre-BAL timepoints (p<0.001). Four hours post-BAL, the CT scores in the left lung where still increased but the difference was not significant anymore (p=0.19). Additionally, post-BAL two animals showed a positive CT score in the right lung but had returned to baseline values at four hours post-BAL. In two timepoints a positive CT score in the right lung was observed at the CT 24 hours post-BAL, the reason for this is unclear.

### Quantitative assessment non-infected group

In Figure 2, the histograms of a representative animal (RM 2) are visualized. These histograms illustrate the relative HU value, and with this density of the lungs, in all timepoints.

**Figure 1:**
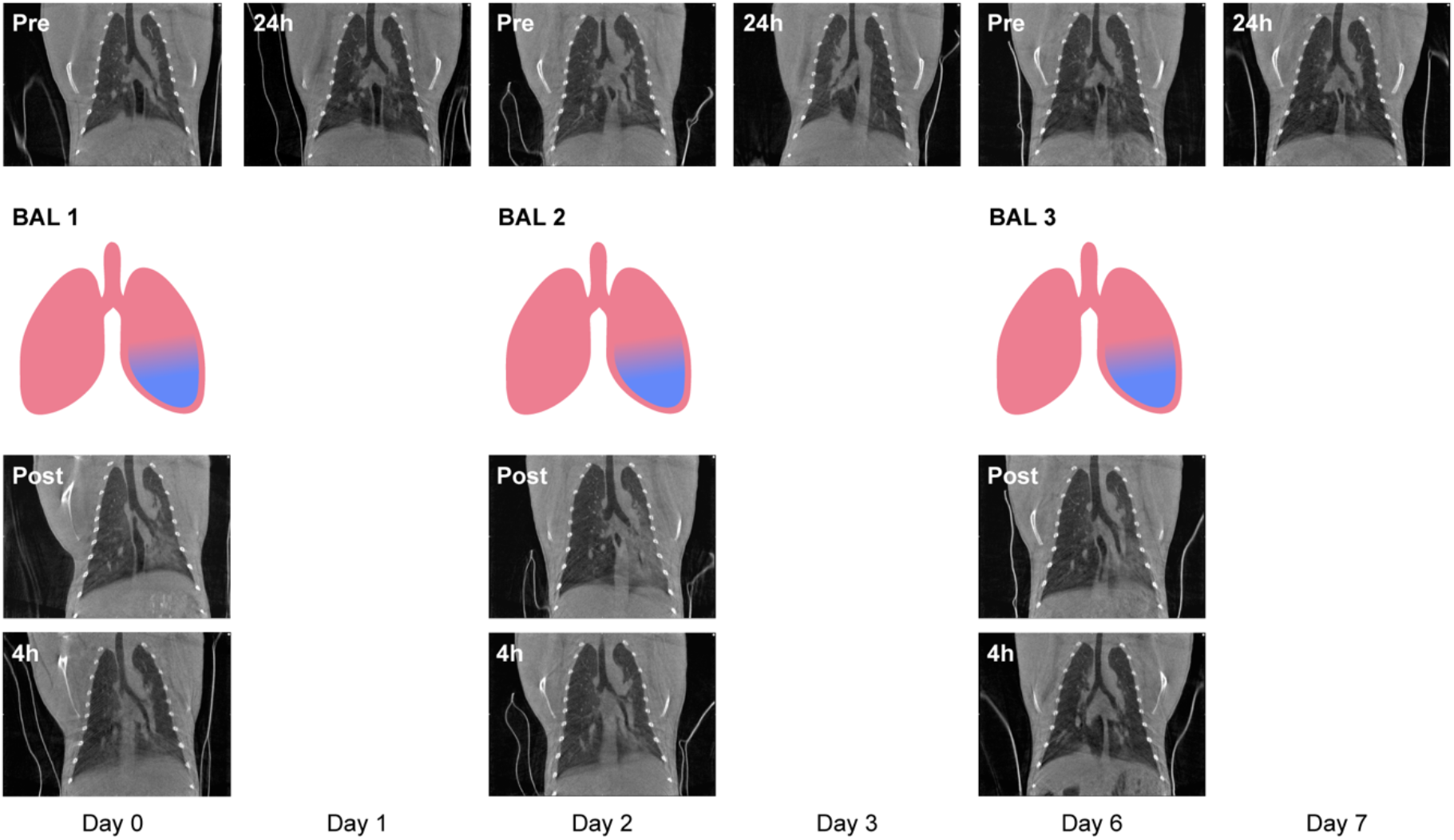
representative coronal slices (of RM2) of all CTs obtained during the three BALs. The procedures are represented in a vertical manner whereas the time in days is represented horizontally. The procedure starts with a pre-BAL CT (Pre), followed by a left-sided (blue) BAL 1, a post-BAL CT (Post) and a four hour post-BAL CT (4h) on Day 0. The next day, Day 1, the 24 hour post-BAL CT (24h) was obtained. The procedures for BAL 2 and BAL 3 were performed in a similar order. Day 4 and Day 5 are not presented in this figure because no procedures were performed on these days.

**Figure 2:**
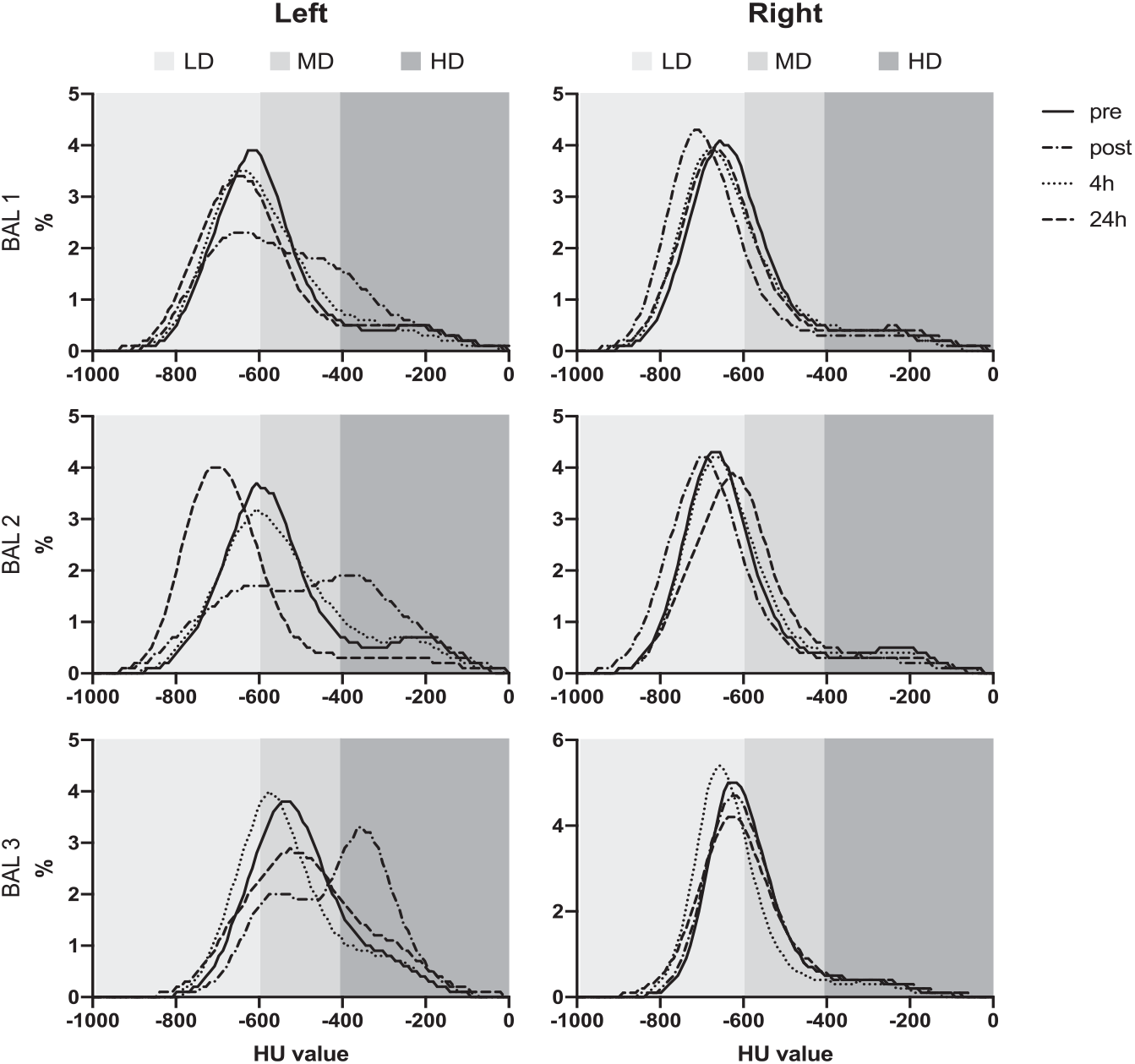
histograms representing the relative lung density pattern of the left and right lung of one representative (RM2) of the non-infected group. LD= −1000/−600 HU, healthy lung, MD= −600/−400 HU, HD= −400/0 HU, non-healthy lung.

There was a significant difference for all three BALs visible at the post-BAL CT for the left lung for both the LD and HD range; LD (decrease in AUC of 30.6%, p=0.003), HD (increase in AUC of 86.7%, p= 0.001). At four hours post-BAL, this difference was still significant compared to the pre-BAL values, now also for the MD range (decrease in AUC of 10.0%, p=0.026), but the AUC change reversed for both LD (increase in AUC of 16.5%, p=0.014) and HD (decrease in AUC of 14.4%, p=0.012) for the left lung. There were also differences found within the right lung, for the four hour timepoint, for the entire histogram; LD (increase in AUC of 11.5%, p=0.012), MD (decrease in AUC of 19.8%, p=0.026), HD (decrease in AUC of 9.4%, p=0.079). For 24 hours post-BAL, no significant differences were observed.

### Visual assessment SARS-CoV-2 infected group

In Figure 3, representative pre-BAL CT and 24 hours post-BAL CT cross sections are shown of two animals, while infected with SARS-CoV-2. The lesions found in the animal visualized in the upper row (RM 13) were minimal and slightly increased on the 24 hours post-BAL timepoint (Video 5-6). For the animal visualized in the bottom row (RM 8), the density of both lungs was increased. For the left lung multiple consolidations were observed on the 24 hours post-BAL CT. On the left side the entire lung, predominantly dorsal, was affected whereas on the pre-BAL CT the majority of the lesions were found in the lower lobe in combination with a ggo in the upper lobe. In addition, in the right lung moderate lesions were detected (Video 7-8).

**Figure 3:**
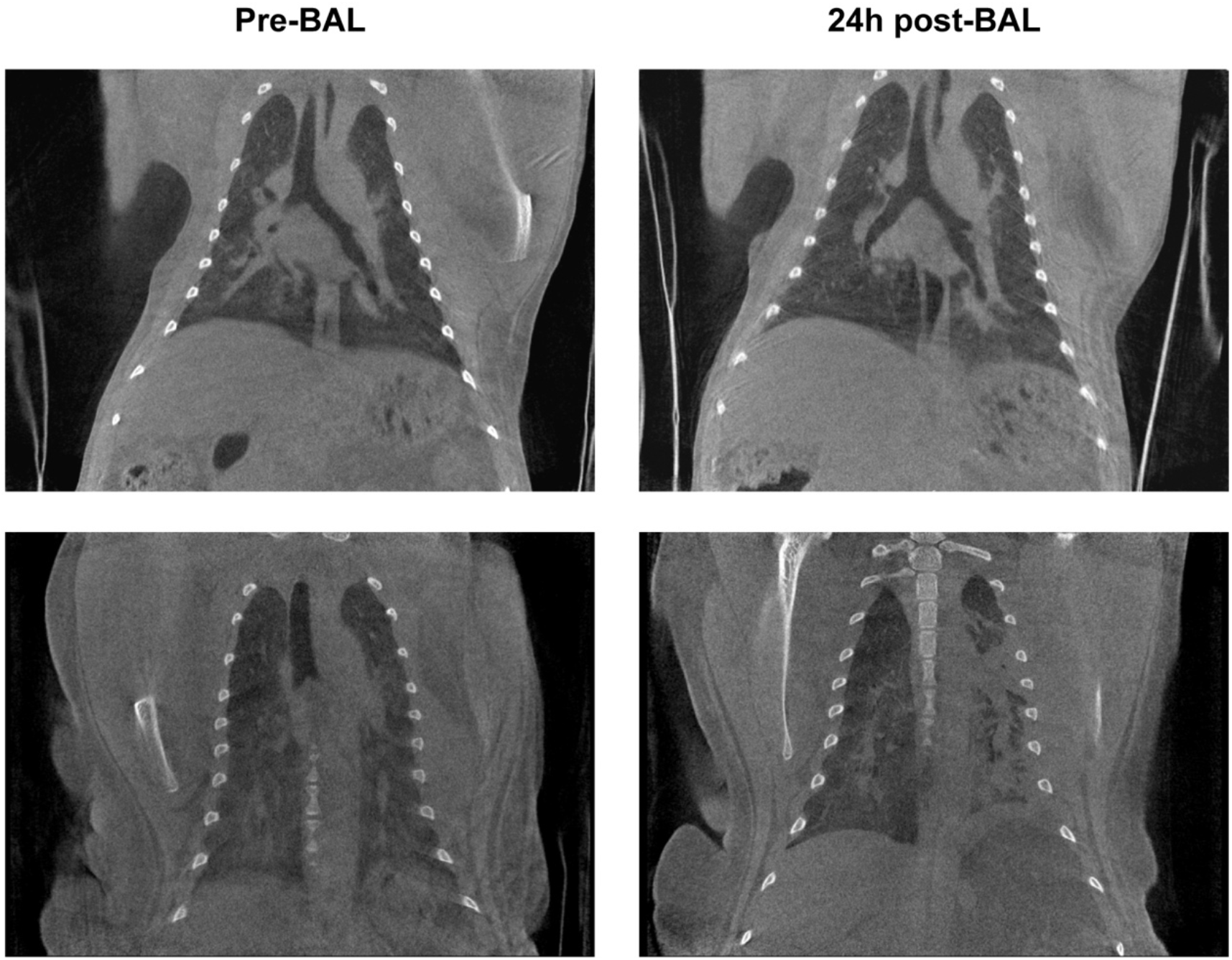
CTs of two infected animals. Upper row RM13; bottom row RM8. Pre-BAL on the left side, and 24 hours post BAL on the right.

In line with the SARS-CoV-2 infection all animals had a positive CT score in at least one side of the lung at the pre-BAL timepoint. In these animals the CT score of the left lung 24 hours post-BAL was significantly increased (p=0.007) compared to pre-BAL CT score with a positive correlation (r=0.907 p<0.0001). For the right lung this difference was not observed. The individual scores of these infected macaques are summarized in Table 2. As previously described a daily CT score of 35 can be reached for the entire lung with a maximum of 15 and 20 for the left and right lung, respectively.

**Table 2:** overview of the CT scores of the left (le) and right (ri) lung of the SARS-CoV-2 infected rhesus macaques

| RM | CT score pre |  | CT score 24h post |  |
| --- | --- | --- | --- | --- |
|  | le | ri | le | ri |
| 5 | 0 | 2 | 0 | 1.5 |
| 6 | 1 | 0 | 1 | 0 |
| 7 | 5 | 0 | 10 | 0 |
| 8 | 6 | 4 | 11 | 6 |
| 9 | 3 | 5 | 6 | 5 |
| 10 | 3 | 1 | 9 | 1 |
| 11 | 0 | 0 | 4 | 0 |
| 12 | 1.5 | 0 | 1 | 0 |
| 13 | 2 | 0 | 4 | 0 |
| 14 | 0 | 5 | 0 | 7 |
| 15 | 1 | 6 | 3 | 2 |

### Quantitative assessment SARS-273 CoV-2 infected group

In Figure 4, the histograms of two representative SARS-CoV-2-infected animals are visualized. These histograms illustrate the relative AUC for the HU values, and with this density of the lungs of the CT obtained before BAL and 24 hours after BAL.

**Figure 4:**
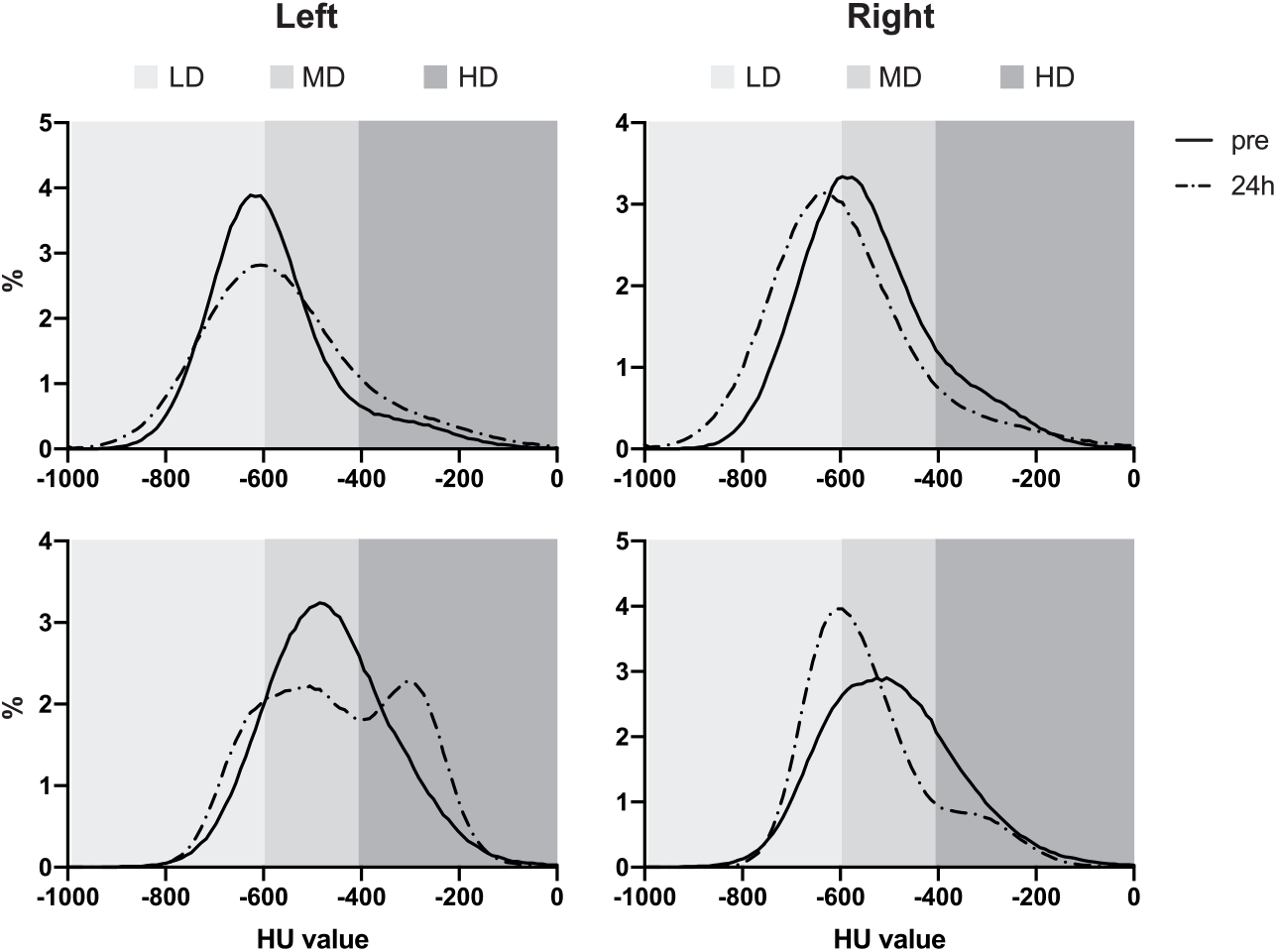
histograms representing the relative lung density pattern of the left and right lung of SARS-CoV-2 infected macaques.

A significant increase in AUC (19.4%) was found for the right lung in the LD range (p=0.004) and a significant decrease in AUC (16.6%) in the MD range (p=0.002). For the left lung a significant difference was only found in the HD range (increase in AUC of 23.2%, p=0.016).

### Comparison BAL 1 versus BAL 2

For investigating the impact of subsequent BALs obtained within a time range of a couple of days, we had seven animals (three non-infected and four infected) from which two BALs were taken within 48 hours. A significant difference in AUC, for the left lung, at the pre-BAL 2 CT was found for both the LD range (decrease in AUC of 27.3%, p=0.022) and MD range (increase in AUC of 18%, p=0.010) and a trend within the HD range (increase in AUC of 55.2%, p=0.076). Also, for the CT score a significant increase was observed for the left lung (p=0.016) at this pre-BAL 2 timepoint compared to the pre-BAL 1 timepoint. To explore whether there are differences in the CT scores between the timepoints in both BALs, the CT scores were subtracted from their respective pre-BAL CT score. In this way, a significant increase in CT score was found for the left lung for the post-BAL timepoint of BAL 1 versus BAL 2 (p=0.047) and for the 24 hours post BAL timepoint of BAL 1 versus BAL 2 (p=0.031). In addition, for these values a positive correlation was found (r=0,77). These differences, for the 24 hours post BAL timepoint, were also detected for all ranges of the histogram; LD (p=0.023), MD (p=0.042) and HD (p=0.015). For the right lung, no significant differences were found for lung density and CT scores or subtracted scores.

**Figure 4:**
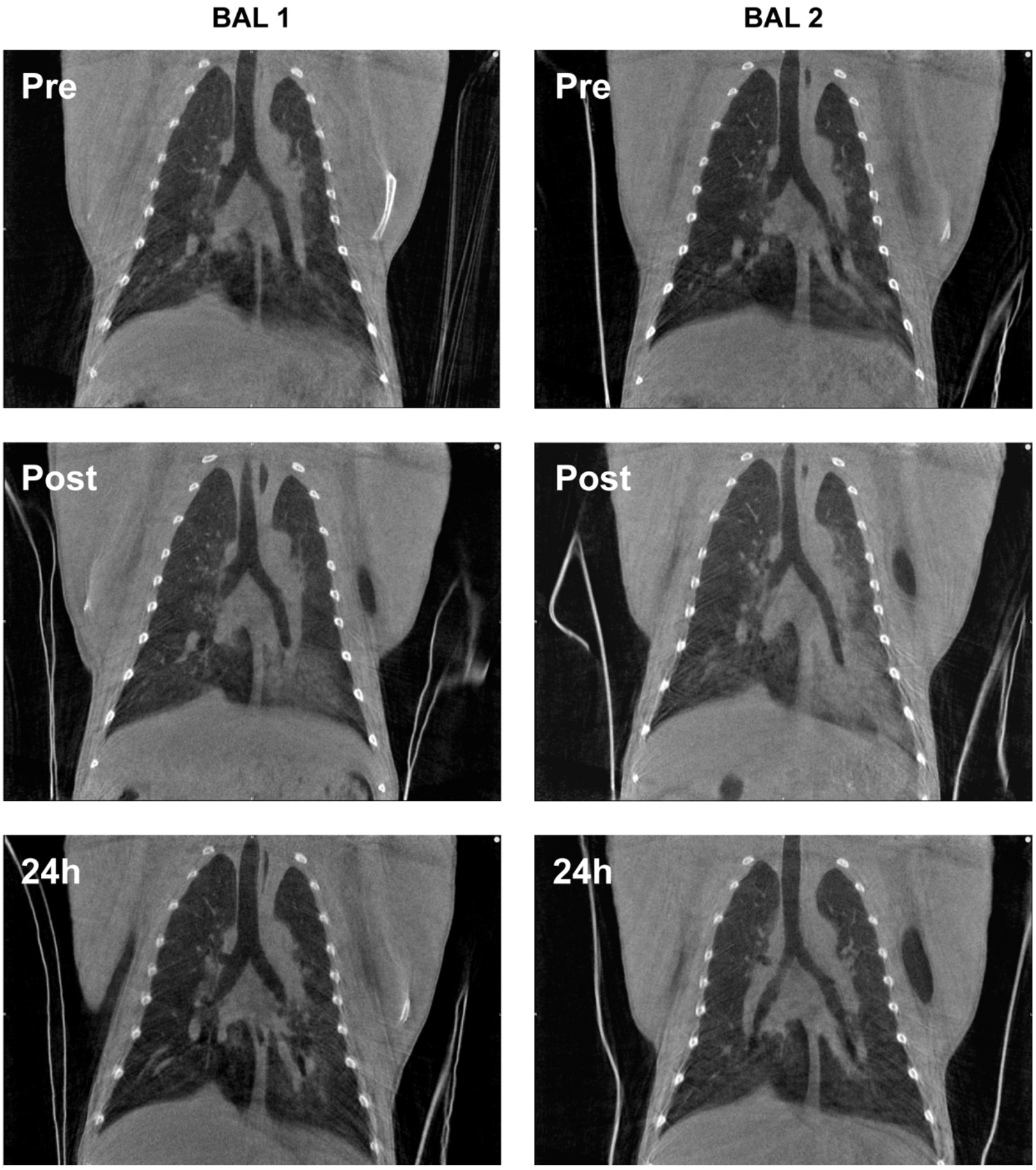
representative coronal CT slices of the first and second BAL of one rhesus macaque

## Discussion

In infectious lung diseases, both CT and BAL are valuable diagnostic modalities which are often executed in preclinical and clinical research. However, the possible interference of BAL on CT has not been investigated so far in NHPs. The findings in this study demonstrate that, in general, thorax CT alterations due to BAL seem to be resolved within 24 hours in non-infected macaques. However, for SARS-CoV-2 infected animals, significant differences for both the semi-quantitative CT score and quantitative AUCs were found at the 24 hours post-BAL CT. In addition, the response of the second BAL within 48 hours of the first BAL shows significant alterations supporting the idea that the effect of the BAL is not finished after 24 hours and therefore might influence acquired CT data.

The pulmonary changes induced by BAL fluid were most pronounced on the immediate post-BAL CT. On those images, increased radiographic opacities were found due to the BAL procedure. The majority of these opacities decreased within four hours after BAL and returned to normal within 24 hours after the procedure. Despite differences in BAL procedure, the changing opacities in time after BAL seen in the macaques are comparable to results obtained in humans (15) and dogs (12). In both studies, a known amount of unrecovered BAL fluid was left in the lung, bilateral in humans and unilateral in dogs. In addition, in dogs no aspiration or repeated instillation was performed. In this study, the procedure was similar to commonly used BAL techniques including repeated fluid instillation and subsequent aspirations. As for the findings in the right lung, the CT score immediately post-BAL was possibly caused by redistribution or a spill of the retained saline due to a change of body posture or by coughing (15). This was supported by the fact that at four hours post-BAL no alterations where found in the right lung anymore. In addition, an explanation for the CT score found 24 hours post-BAL remains unclear.

The current study does not provide evidence of an increased percentage of retained fluid but redistribution between the lung lobes due to coughing cannot be excluded. The differences found between BAL 1 and BAL 2 suggest that the effect of these procedures is still ongoing 24 hours after subsequent BALs, however not clearly visible. As BAL 3 was obtained four days after BAL 2, it was not expected to find ongoing effects that could be related to BAL 2. By comparing all the outcome parameters from BAL 1 with BAL 3 and BAL 2 with BAL 3 significant differences were not found.

Multiple publications showed that NHPs are susceptible to SARS-CoV-2 infection and develop a mild-to-moderate form of disease (5–8), with the discrimination made between mild and moderate based on the presence or absence of imaging findings (26). In the current study, the majority of the infected animals (15/17) showed CT scores above zero at the pre-BAL timepoint. In addition, infection with SARS-CoV-2 was confirmed in each animal by RT-PCR. The strong positive correlation that was found between the pre-BAL and 24 hour post-BAL CT scores indicates that a BAL can potentially enhance the visual pathology of infected lung tissue. As COVID-19 is a rapidly developing infection it cannot be excluded that the increase in CT score is due to a worsening of the disease. However, in patients fluctuations in thorax CT abnormalities are not observed during the active phase of the infection. The active phase can be divided in four stages; early (day 0-5), progressive (day 5-8), peak (day 9-13) and late stage (> 14 days) suggesting a Gaussian curve in the temporal evolution of lung abnormalities (27). This is supported by the observation during multiple studies at our facility, that SARS-CoV-2 infected rhesus macaques show an equal or even lower score in subsequent CTs obtained over 24 hours after BAL though before ten days post infection (data not shown). The current study suggests that these fluctuations found are induced by the BAL as all scans are obtained within the first ten days of infection.

Furthermore, the right lung showed less lesions compared to the left lung. As the number of lesions reflected by the CT score seems to influence the outcome this may be of impact. Since the infection with SARS-CoV-19 went by intratracheal and intranasal route it was expected that both lungs would reveal a similar CT score. However, in the animals included in this study, the left lung showed more lesions compared to the right lung. An unexpected finding, as this was not observed in other SARS-CoV-2 infection studies performed at our institute. What was observed is the high level of heterogeneity between the animals in the infectious take of SARS-CoV-2. This is probably coincidence-as our animal selection was solely based on availability of BALs coupled with CTs.

As expected the animals showed an increase in HD range of the left lung, caused by the remnants of the BAL fluid. More interestingly, an increase in the LD range in their non-lavaged right lung was found. Amigoni *et al*. showed in mice, that the relative tidal volume of the contralateral lung of an injured lung increased while mechanical ventilated (28). These data support the hypothesis that the contralateral lung will develop a coping mechanism to ensure sufficient oxygenation. The AUC decrease in LD and MD versus increase in HD immediate post-BAL can be explained by the instillation of relative HD saline. These findings are in corroboration with the measured post-BAL non-air volumes in humans (15). At four hours post-BAL, the reversed significant difference, compared to the pre-BAL values for both LD and HD for the left lung, could probably be explained as a similar compensating reaction described above to the initial loss of LD tissue in the lavaged lobe of the lung. This assumption can neither be confirmed nor completely denied with the limited amount of data available on this topic. Further research on lobar regions with more subjects is necessary to confirm these findings.

Another objective was to investigate the impact of multiple BALs on CT. Therefore, we compared BAL 1 and BAL 2, performed within 48 hours in seven animals. It was interesting to find a difference in AUC, for the left lung, in all ranges. These findings confirm the impact of the BAL on the recovery time based on the readout of thorax CTs when multiple BALs are performed in a short period of time (13). As four animals in this group were infected, although unlikely, artificial dissemination of SARS-CoV-2 cannot be excluded.

Whereas the discussion concerning the diagnostic value of both BAL and CT in human is ongoing (15, 18), there is consensus of the value of both procedures in COVID-19 NHP models (6, 13, 14, 19). The current study shows that when using both BAL and CT it is of importance to acknowledge the existing interference between those two valuable techniques in rhesus macaques.

## Author contributions

Conceptualization, AM, BV, EV, MS; methodology, AM, LM, JB, LG, MS; formal analysis, AM, LM, DZ, MS; investigation, AM, LM, MS; writing-original draft preparation, AM and MS; writing – review and editing, AM, LM, JB, DZ, JL, EV and MS; supervision; JL, MS. All authors have read and agreed to the published version of the manuscript.

The authors declare no conflict of interest.

## Acknowledgements

We would like to thank the Animal Science Department, especially the animal caretakers for excellent care of the animals and Francisca van Hassel for her assistance with figure editing. Part of this work was supported by internal funding from the Biomedical Primate Research Centre.

## Notes

### Competing Interest Statement

The authors have declared no competing interest.

